# The impact of human agents on spatial navigation and knowledge acquisition in a Virtual Environment

**DOI:** 10.1101/2024.12.18.628314

**Authors:** T. Sánchez-Pacheco, M. Sarría-Mosquera, K. Gärtner, V. Schmidt, D. Nolte, S.U. König, G. Pipa, P. König

**Affiliations:** Institute of Cognitive Science, University of Osnabrück, Wachsbleiche 27, 49090 Osnabrück, Germany; Department of Neurophysiology and Pathophysiology, University Medical Center Hamburg-Eppendorf, Martinistr. 52, 20246 Hamburg, Germany

**Keywords:** Spatial Navigation, Human Agents, Virtual reality, Exploration-Exploitation, Social Facilitation

## Abstract

Concepts of spatial navigation rest on the idea of landmarks, which are immobile features or objects in the environment. However, behaviorally relevant objects or fellow humans are often mobile. This raises the question of how the presence of human agents influences spatial exploration and knowledge acquisition. Here, we investigate exploration and performance in subsequent spatial tasks within a virtual environment containing numerous human avatars. In the exploration phase, agents had a locally limited effect on navigation. They prompted participants to revisit locations with agents during their initial exploration without significantly altering overall exploration patterns or the extent of the area covered. However, agents and buildings competed for visual attention. When spatial recall was tested, pointing accuracy toward buildings improved when participants directed their attention to the buildings and nearby agents. In contrast, pointing accuracy for agents showed weaker performance and did not benefit from visual attention directed toward the adjacent building. Active agents and incongruent agent-environment pairings further enhanced pointing accuracy, revealing that violations of expectations by agents can significantly shape navigational knowledge acquisition. Overall, agents influenced spatial exploration by directing attention locally, with the interaction between agent salience and environmental features playing a key role in shaping navigational knowledge acquisition.

## 1 INTRODUCTION

Spatial navigation is essential for goal-oriented movement and active environmental interaction (Epstein et al., 2017; Ito et al., 2015). In humans, regular engagement in spatial navigation, whether studied through navigation done in the context of professional activities (Griesbauer et al., 2022; Maguire et al., 2006; Woollett and Maguire, 2011), targeted training (Choi et al., 2012), or virtual environments (West et al., 2017), is related to the enhancement of cognitive functions, particularly memory and spatial awareness. This intricate link between spatial navigation and cognitive processes highlights the crucial role of spatial navigation in developing and organizing spatial knowledge.

The transformation of navigational experiences into spatial knowledge begins with recognizing key environmental elements, followed by their gradual integration into a cohesive reference system (Ekstrom and Isham, 2017). When individuals explore new surroundings freely, they can identify elements that aid their orientation and harness them as building blocks for knowledge construction, such as remembering shops when visiting a new town. This process, known as landmark identification, is essential for maintaining positional awareness and planning future pathways (Janzen et al., 2006). Spatial knowledge encompasses recognizing and recalling critical elements of the landscape, understanding their spatial relationships, and the routes connecting them.

Despite extensive research on the role of static landmarks in spatial cognition (Malanchini et al., 2020), the dynamic human aspects influencing spatial exploration and knowledge acquisition have yet to be fully explored. Often, humans are viewed primarily as modifiers of navigational paths rather than as active contributors to spatial knowledge acquisition (Bicanski and Burgess, 2020; Ekstrom and Isham, 2017). This perspective overlooks the significant role other individuals play in real-world spatial cognition. Encounters with others can impact pedestrian dynamics (Dalton et al., 2019), influence visual exploration (Gert et al., 2020), provide vital information about the safety and usability of spaces (Bajorunaite et al., 2022), affect the recall of locations (Kuehn et al., 2018), and prompt a parallel social mapping of the environment (Schafer and Schiller, 2018). Thus, incorporating other humans into navigation research is essential for a deeper understanding of spatial cognition.

However, studying the influence of fellow humans on concrete spatial knowledge faces the difficulties of maintaining a navigation scenario of real-world scale in controlled environments. When available, researchers often conduct retrospective studies using real-world data, such as mobile phone and GPS data, which can characterize human mobility patterns but lack the controlled variables necessary for isolating specific factors mediating the results (Pappalardo et al., 2015; Schläpfer et al., 2021). Moreover, these studies do not link exploration patterns to the future acquisition of spatial knowledge, hindering a comprehensive understanding of cognitive processes derived from navigation patterns. When moving to the other side of the spectrum, laboratory setups tend to be bound to small-scale environments and simplified tasks that lack the challenge of spatial scales of the real world (Wiener et al., 2020). Participants are asked to use humans as ankers inside a maze-like VR or as landmarks in visual flow experiments (Kuehn et al., 2018). Dalton et al. (2019) point out that VR technologies in wayfinding research often exclude the presence of others, thereby underestimating their importance. They suggest that the mere presence of others (i.e., weak social cues), even without active interaction, can help infer the importance of a space within its environment, and actively call for more studies to be done in this realm of research. Hence, it is acknowledged that examining how the mere presence of others might impact navigation and knowledge acquisition when humans are not primary targets is essential but remains unstudied.

With this work, we raise the question of whether spatially relevant information can be extracted from observing other humans in the space around us. It is important to note that humans do not inherently serve as fixed landmarks in natural navigation, primarily due to their mobility, preventing them from forming intrinsic associations with specific places. However, rather than considering other humans as reliable reference points, we explore the notion that their relevance within a given context can be harnessed as a source of information to enhance spatial knowledge. Essentially, it is not the presence of others per se that aids in spatial cognition but rather how they interact with the elements in their surroundings that could be relevant for spatial exploration and knowledge acquisition. This prompts us to investigate what minimal change in a human agent’s interaction with the environment can elicit participants’ different behavioral responses. We seek to understand whether these human elements act as distractions, potentially diminishing the saliency of the surrounding stimuli, or if, conversely, they contribute to improved performance by anchoring individuals to a more vivid mental representation of specific locations.

To address this complexity, we investigated how spatial knowledge acquisition developed in a controlled one-square-kilometer virtual reality (VR) environment with human agents. The study incorporated human agents at two levels: one in which the agent interacted with the environment by holding an object relevant to the context, such as a toolbox in front of a hardware store (i.e., Active agent), and another in which the agent simply stood without interacting with any objects around it (i.e., Passive agent). These agents were placed in front of public buildings, such as stores, basketball courts, restaurants, or residential buildings. For our first group of participants, all active agents were placed in front of public buildings that matched their object interaction. For the second group, we disrupted this congruency to study the sole influence of the type of agent and their building on the context in which they are situated, under the hypothesis that having context-congruent agents will enhance the participant’s ability to recall. The aim of our study is to explore the role of these human agents and their congruency with context in spatial exploration and the development of spatial knowledge.

## 2 RESULTS

We examined the impact of human agents on spatial navigation and knowledge acquisition in a virtual city named Westbrook, consisting of 236 buildings. We identified 26 public buildings (e.g., shops, basketball court) and 26 residential buildings as task-relevant, marking them with street art. Additionally, there were 180 buildings without graffiti and four large buildings on the city’s outskirts, which could serve as global landmarks given their dimensions. We designed two categories of human agents: active agents, who performed context-relevant actions (e.g., holding a toolbox in front of a hardware store), and passive agents, who held a resting position without interacting with objects. In the first experiment, active agents were placed in public areas, displaying actions congruent with the buildings, while passive agents were positioned in front of residential buildings without interaction. In the second experiment, both agent types were split evenly across public and residential areas, disrupting the congruent pairs. Participants in both experiments completed five 30-minute exploration sessions, totaling 150 minutes. Additionally, to provide a baseline for comparison of the exploration strategies, we used the control group from Schmidt et al. (2023) who explored the same VR city (Westbrook) with the same session lengths and numbers but no agents present. We investigated the exploration phase by analyzing participants’ navigational coverage of the city, their walking strategies, agent-induced bias in their exploration, and their visual behavior during exploration. Finally, we tested their spatial knowledge acquisition in a separate session using VR pointing tasks.

To establish comparability in spatial orientation abilities between the participants of the experiments (experiment 1, experiment 2, and control), participants completed the FRS (Fragebogen Räumlicher Strategien) questionnaire, and their scores were contrasted. There were no significant differences between the groups at baseline on any of the three subscales (global, survey, and cardinal), *χ*^2^(2, *N* = 67) ≤ 1.68, *p* ≥ 0.43. Therefore, the groups were comparable in their assessment of their use of spatial strategies before the start of the experiments.

### 2.1 Assessment of the exploration phase

During the VR city experiment, we tracked participants’ exploration, including walking behavior, navigational coverage, decision-point strategies, and visual behavior. This comprehensive analysis revealed their navigational strategies and engagement, highlighting the agents’ impact on their exploration.

#### 2.1.1 Navigational Coverage of the City

We quantified participants’ walking behavior in the virtual city using a primal city graph (Neal, 2013) to analyze participants’ free exploration patterns. In this graph, decision points (i.e., intersections of walkable paths) were represented as nodes and paths connecting them as edges. Participants’ navigational coordinates (see Figure.2a) were assigned to the nearest graph element (see Figure.2b), defining their exploration as movements from one graph element to another. Out of 159 nodes, participants visited between 45 and 113 unique nodes during each 30-minute session (*M* = 87.06, *SD* = 16.49). We calculated the coverage ratio by dividing the number of nodes visited at least once by the total number of nodes.

**Figure 1.**
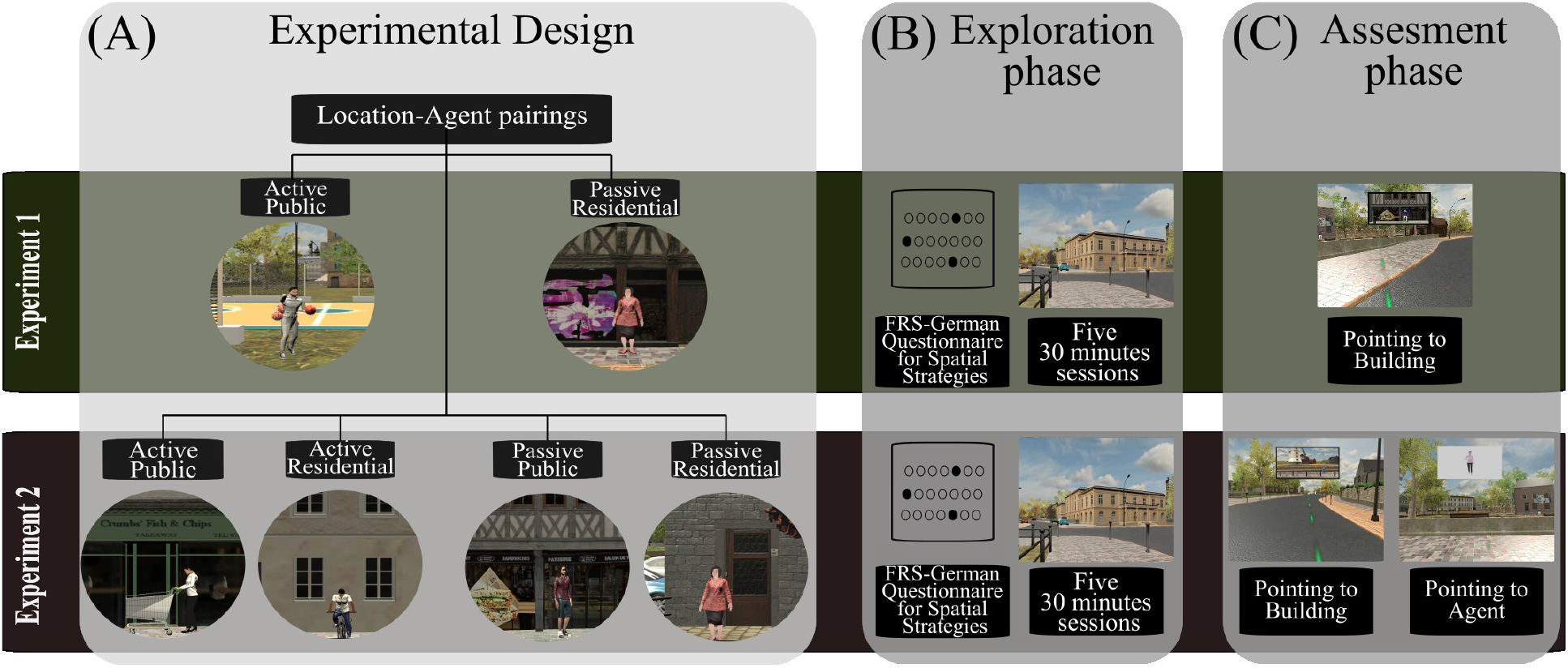
Experimental design and Procedure: Panel A illustrates our 2×2 experimental design, featuring Passive/Active agent types and Public/Residential buildings, which were assigned differently in the two experiments. Panel B outlines the exploration phase procedure, where participants first filled out the FRS questionnaire and then completed five 30-minute sessions of unguided exploration within the virtual city of Westbrook. Panel C describes the assessment phase during a sixth session, where participants completed pointing tasks within the VR city.

**Figure 2.**
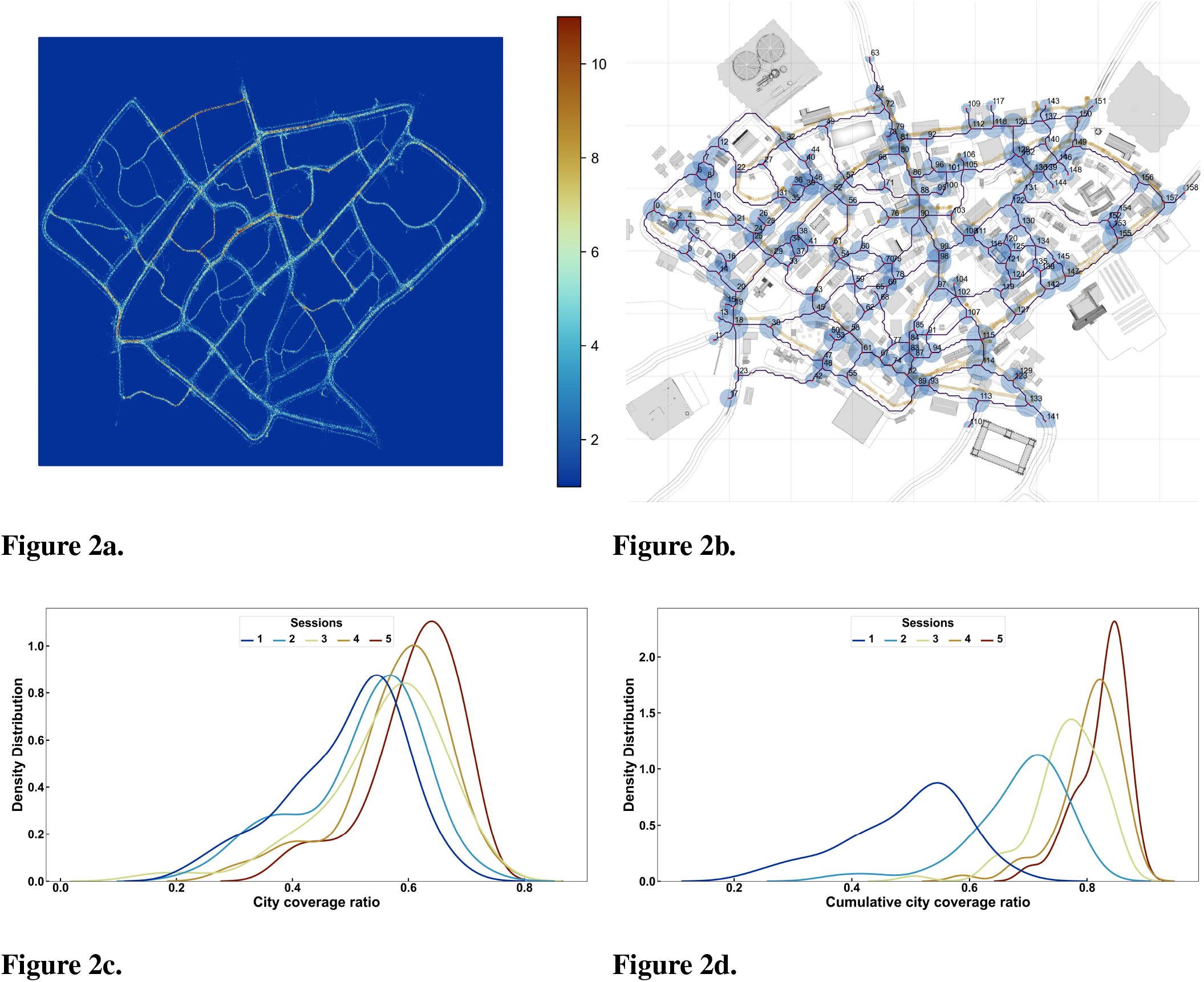
Free exploration of Westbrook: **(A)** presents a heatmap that visualizes the spatial navigation data of all participants across the exploration phase. It compiles movement across all sessions, with participant locations discretized to the nearest centimeter and averaged on a second-by-second basis after removing the initial two seconds of each recording. We fitted a heatmap grid approximating a 1-centimeter resolution and created a color scale ranging from 0 to 10 visits, highlighting areas of varied exploration intensity. **(B)** shows Westbrook from above, overlaid with a graph structure we used to formalize navigational data as decision units. **(C)** quantifies exploration through a map coverage ratio, calculated by dividing the number of unique nodes visited by the total nodes in the city. **(D)** compiles these ratios on a cumulative basis, reflecting the participants’ expanding discovery of the city over time.

To account for repeated measures, we implemented a linear mixed-effects model (LMM) that considered intrasubject variability and generated individual intercepts for each participant. This model predicted the individual session coverage ratio as a function of the session, the experimental group (control versus city with agents), and their interaction. Using the first session as a baseline, we observed significant cumulative increases in navigational coverage with each subsequent session. Starting from an individual coverage of roughly half the city (as shown by the mean of the blue curves in Figure.2c and Figure.2d), the LMM analysis demonstrated that session effects progressively increased the individual coverage ratio. The coefficients indicate changes relative to session 1, with positive values signifying an increase: *β*_Session 2_ = 0.03 (SE = 0.01, *t* = 2.48, *p* = 0.01), *β*_Session 3_ = 0.07 (SE = 0.01, *t* = 6.54, *p <* .001), *β*_Session 4_ = 0.09 (SE = 0.01, *t* = 8.53, *p <* .001), and *β*_Session 5_ = 0.12 (SE = 0.01, *t* = 11.49, *p <* .001).

The effect of the experiment group (agents versus control) on the coverage ratio was not statistically significant (*β*_Experiment_ = −0.02, SE = 0.02, *t* = −0.72, *p* = 0.48). Additionally, interactions between the session and experiment group were also not significant: *β*_Session i:Experiment_ ≤ − 0.04, *p* ≥ .07. This demonstrated that as sessions advanced, participants covered more ground within the same timeframe, achieving this equally in both a city with agents and one devoid of them.

In order to estimate if the participants were deferentially accumulating unique decision points in the city as the sessions went on, we calculated a cumulative coverage ratio, accumulating the number of uniquely visited nodes as the sessions progressed. A linear mixed-effects model (LMM) with participants as random effects was used to predict the cumulative coverage ratio as a function of session, experiment group, and their interaction. This analysis showed a significant progression, with participants covering more of the city in each subsequent session (see Figure.2d). Starting from the same initial coverage ratio mean (*M* = 0.49), significant cumulative increments in navigational coverage were observed in each session: *β*_Session 2_ = 0.19 (SE = 0.01, *t* = 24.71, *p <* .001), *β*_Session 3_ = 0.27 (SE = 0.01, *t* = 35.80, *p <* .001), *β*_Session 4_ = 0.31 (SE = 0.01, *t* = 40.75, *p <* .001), and *β*_Session 5_ = 0.33 (SE = 0.01, *t* = 43.48, *p <* .001). The effect of the experiment group on the coverage ratio was not statistically significant (*β*_Experiment_ = −0.02, *p* = 0.48). Additionally, interactions between the session and experiment group were also not significant: *β*_Session i:Experiment_ ≤0.02, *p* ≥ .33. After the fifth session, the end of the exploration, the cumulative number of unique nodes visited (*M* = 0.83, *SD* = 0.04) indicated that most participants had seen the majority of the city. These findings suggest that the presence of an agent did not significantly influence the navigational coverage of the city.

#### 2.1.2 Exploration strategies on decision points: Exploratory vs. conservative behavior

To investigate participants’ use of conservative or exploratory walking strategies, we analyzed whether the picked to walk towards had been visited more often than the comparative directions (conservative) or if they preferred the less frequented paths (explorative). We defined discrete navigational decisions as movements from one node to another and quantified them using a strategy matrix (see Figure.3a and 3b). A linear mixed-effects model was fitted to the data to examine differences in decision numbers based on session, strategy (conservative vs. explorative), and experiment (control vs. city with agents), with random intercepts for participants to account for the nested data structure. The model’s fixed effects indicated that the average number of decisions across all factors was *β*_0_ = 73.14 (SE = 4.02, *t* = 18.20, *p <* .001). The analysis showed significant increases in decisions as participants gained experience in Westbrook, with increases evident from the first session onward: *β*_Session 2_ = 30.17 (SE = 2.49, *t* = 12.11, *p <* .001), *β*_Session 3_ = 51.23 (SE = 2.49, *t* = 20.57, *p <* .001), *β*_Session 4_ = 67.49 (SE = 2.49, *t* = 27.09, *p <* .001), *β*_Session 5_ = 80.32 (SE = 2.49, *t* = 32.24, *p <* .001). The difference in decisions between conservative and exploratory strategies was significant, *β* = 75.52 (SE = 3.52, *t* = 21.44, *p <* .001). The interaction between session and strategy was significant, showing that decision increases per session were lower for the exploratory strategy compared to the conservative one: *β*_Session 2:Strategy1_ = −44.70 (SE = 4.98, *t* = −8.97, *p <* .001), *β*_Session 3:Strategy1_ = −67.66 (SE = 4.98, *t* = −13.58, *p <* .001), *β*_Session 4:Strategy1_ = −86.02 (SE = 4.98, *t* = −17.26, *p <* .001), *β*_Session 5:Strategy1_ = −102.37 (SE = 4.98, *t* = −20.55, *p <* .001). No significant difference in decision numbers was observed between the two experiments (*β*_Experiment_ = −3.95, SE = 7.40, *t* = −0.53, *p* = 0.84), indicating that the presence or absence of agents did not markedly influence decision-making strategies. Both behaviors increased as sessions progressed. Initially, participants prioritized exploratory behavior, but as they gained experience, they integrated conservative behavior, effectively combining both strategies (see Figure.3c), regardless of agent presence. This strategy adaptation, where exploratory decisions remained stable while conservative decisions increased, occurred without significant influence from agents, suggesting that the global exploration strategy is unaffected by human agents.

**Figure 3.**
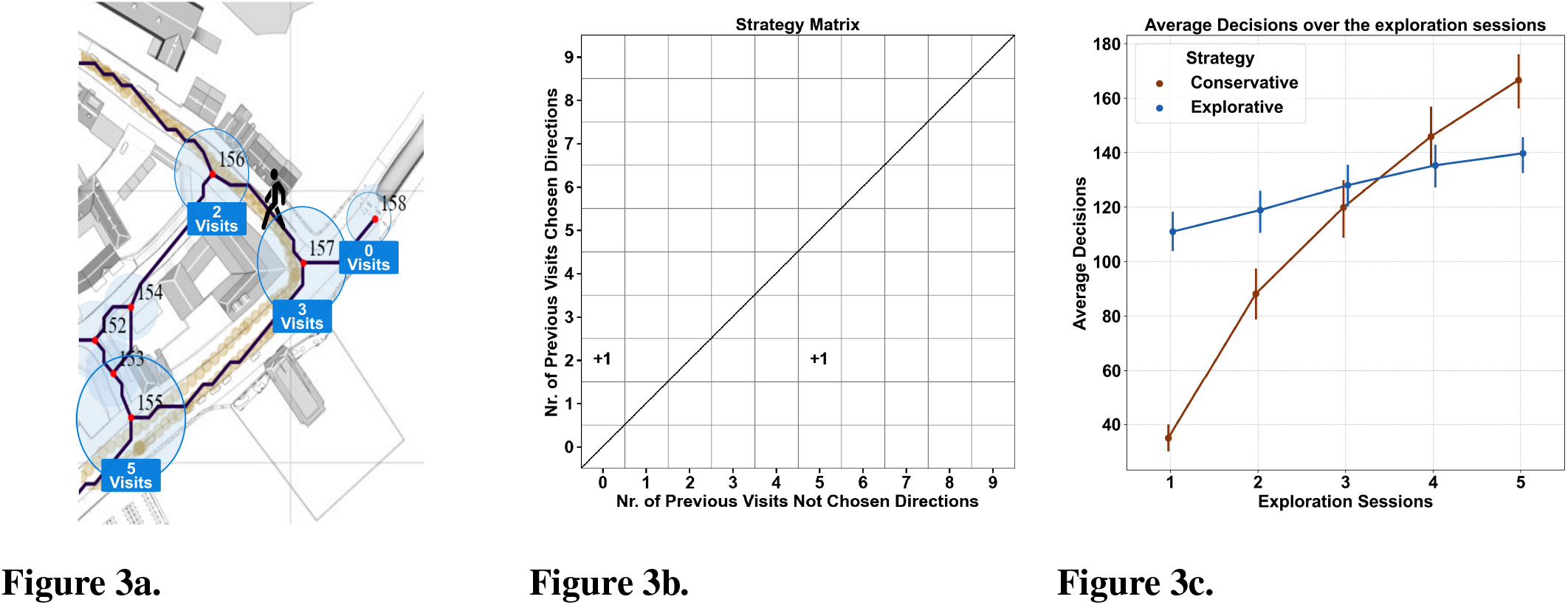
Exploration strategy quantification: **(A)** This panel illustrates the translation of continuous movement into quantifiable behavioral units. Movement coordinates, denoted by beige dots, are assigned to the closest node or edge centroid. Nodes, represented by blue circles, are positioned at decision points and are interconnected by black lines (edges). A *conservative* decision occurs when participants return to a frequently visited node, whereas an *explorative* decision is made when a less frequented node is chosen. **(B)** This panel introduces a strategy matrix designed to analyze navigational choices. The matrix evaluates the path chosen by a participant (displayed on the y-axis) against all other possible paths at a decision point (x-axis). For instance, upon exiting node 157 (as seen on Panel A), the participant’s choice of node 156 (with two previous visits) is compared against nodes 155 and 158, which had 5 and 0 visits, respectively. The matrix is structured such that rows indicate visits to the selected node (in this case, 156) and columns to the unselected nodes (in this case, 155 and 158). Entries in the matrix at positions [2,0] and [2,5] increment by one, reflecting the selection of node 156 over alternatives. This matrix visually encodes decision-making patterns: counts above the diagonal suggest conservative decisions, and those below indicate exploratory actions. **(C)** This panel synthesizes decision-making trends by displaying the average sum of decisions, categorized by strategy, by all participants across our two experiments and the control group in a single session.

#### 2.1.3 Agent-induced bias on walking strategies

To evaluate the impact of agents on participants’ exploratory behavior, we analyzed decision points where participants could choose between a path with an agent and one without, assuming that agents might induce elicit visits. Data from these points were compared with a control group from Schmidt et al. (2023), who explored the same VR city (Westbrook) without agents (See Figure.4a). We used identical decision points in both scenarios. We applied a linear mixed-effects model to assess visit counts, accounting for session and experiment type (Control vs. City with Agents) with random intercepts for participants. Results indicated a significant increase in visit counts across sessions compared to Session 1. Specifically, visit counts in Session 2 were, on average 14.84 points higher (*β*_Session 2_ = 14.84, *p <* 0.001), and this effect continued to grow in subsequent sessions, culminating in a 38.55 point increase by Session 5 (*β*_Session 5_ = 38.55, *p <* 0.001). However, the overall difference between the experiment types was not significant (*β*_Experiment_ = −2.15, *p* = 0.50), suggesting that the presence of agents did not universally affect visit counts across all sessions. Notably, the interaction between session and experiment type was significant in Session 5 (*β*_Session 5:Experiment_ = 5.47, *p* = 0.008), showing a greater increase in visits in the City with Agents group. To further explore the influence of agents, we calculated the likelihood of participants adopting exploratory versus conservative strategies. This was done by dividing the number of choices for each strategy cell (i.e., above and below the diagonal; see Figure.4b) by the total number of decisions made, as recorded in the mirrored cells of the strategy matrix. For instance, we summed the number of times a participant chose to move to a place they had visited only once over a place they hadn’t visited (cell [1,0], conservative behavior) with the number of times they chose to go to a place they had never been over a place they had visited once (cell [0,1], explorative behavior), and divided each count by that total of decisions made in both cells (sum of the count in both [1,0] and [0,1] cells). This indicator showed a clear distinction between the behavior of participants in the city with agents and that of the control group. Within the first exploration session, participants in the city with agents had an average proportion of conservative behavior of *M* = 0.47 (*SD* = 0.16) compared to *M* = 0.20 (*SD* = 0.09) in the control data, while the explorative behavior was *M* = 0.53 (*SD* = 0.16) for the city with agents and *M* = 0.80 (*SD* = 0.09) for the control data. This indicates that agents prompt more local conservative behavior, reducing exploratory actions during participants’ initial city exposure.

**Figure 4.**
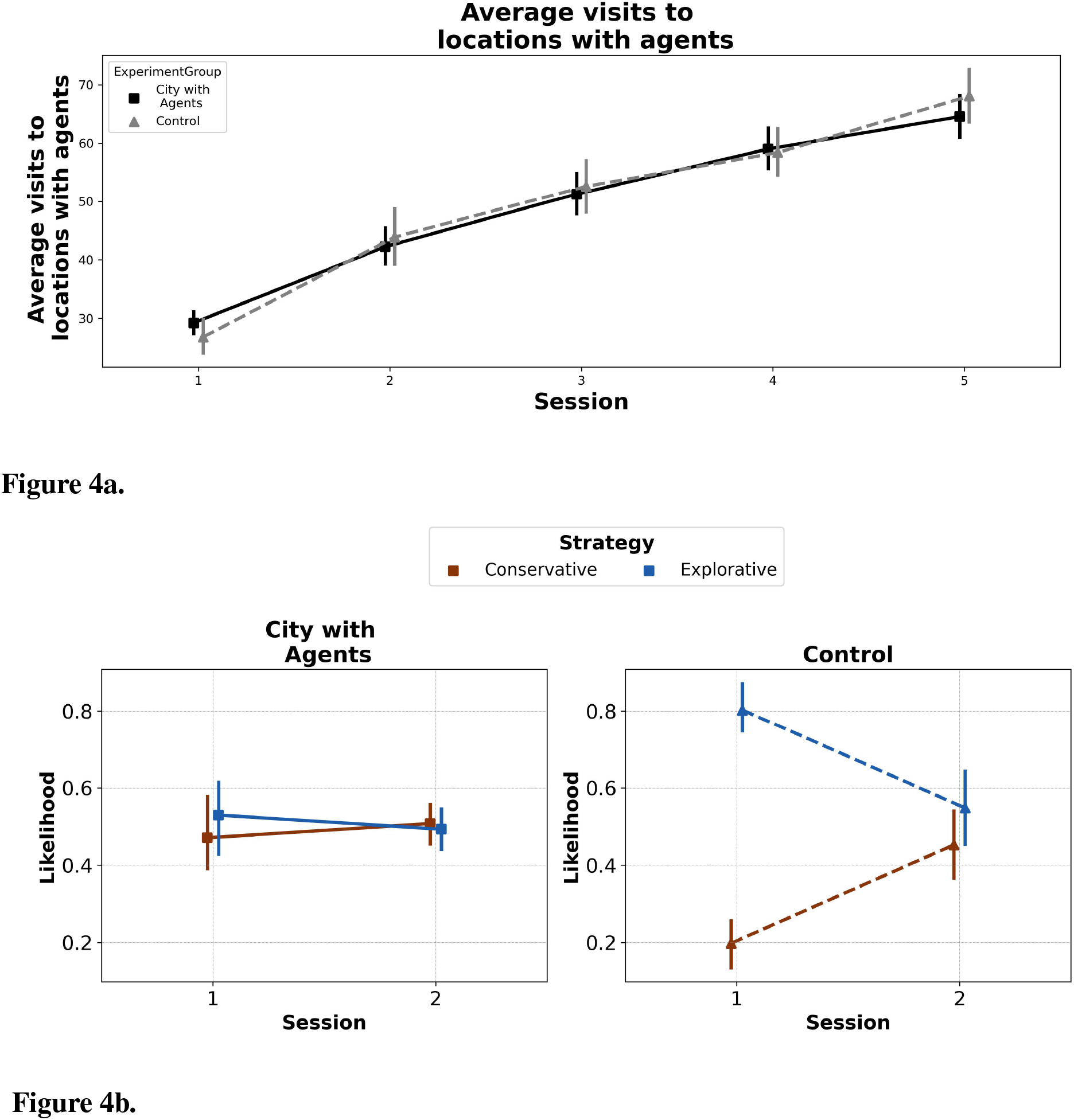
Agent-induced bias in exploration patterns: **(A)** The figure compares decision-making in virtual environments with and without agent presence. It contrasts our experimental data against a control from Schmidt et al. (2021). Panel A shows the average visit count to agent locations in our experiment (black) and the visits to those same points in the control (grey). **(B)** The probability of participants opting for paths where agents are located, classifying decisions as *conservative* when going towards the agent was the more familiar path, and *exploratory* when choosing the agent path was the less frequented route. We only show the first two sessions as the difference disappears afterward.

#### 2.1.4 Assessment of visual behavior during exploration: investigating dwelling time on agents and buildings

To characterize what participants focused on in the city, we quantified their visual behavior by summing the cumulative time spent gazing at each object, termed dwelling time. We hypothesized that participants would have higher dwelling times for active agents and public buildings compared to passive agents and residential buildings, assuming active agents congruent with their surroundings would attract the most attention.

The data revealed distinct patterns in the attention participants allocated to different types of objects in the city. Notably, general residential houses across the city were observed for a shorter duration compared to our experimental buildings, with participants spending on average approximately 6.68 seconds (*M* = 6.68, *SD* = 6.69) gazing at these buildings. This is in contrast to the longer viewing times for our public task buildings (*M* = 11.82, *SD* = 8.48) and residential task buildings (*M* = 11.66, *SD* = 8.29), which attracted more sustained attention from participants. Furthermore, global landmarks within the city garnered significantly longer dwelling times, with participants spending an average of 16.27 seconds (*M* = 16.27, *SD* = 10.69) focusing on these prominent features, nearly three times the average dwelling time of general buildings. In terms of agents, active agents were observed for an average of 3.63 seconds (*M* = 3.63, *SD* = 3.74), while passive agents attracted slightly less attention, with an average dwelling time of 2.66 seconds (*M* = 2.66, *SD* = 2.78). These findings indicate that our experimental manipulations effectively captured participants’ gaze in the expected order for both buildings and agents.

We further analyzed how each experimental factor influenced visual attention. On average, participants spent more time looking at active agents (*M* = 3.39, *SD* = 3.49) compared to passive agents (*M* = 2.73, *SD* = 2.91, see Figure.5a). Active agents also seemed to distract participants from focusing on the area behind them, as buildings with active agents had lower dwelling times (*M* = 12.42, *SD* = 8.13, see Figure.5e) compared to those with passive agents (*M* = 13.28, *SD* = 9.27). These results underscore the adversarial relationship between agents and building attention, where increased focus on active agents corresponded with decreased attention to the buildings behind them.

**Figure 5.**
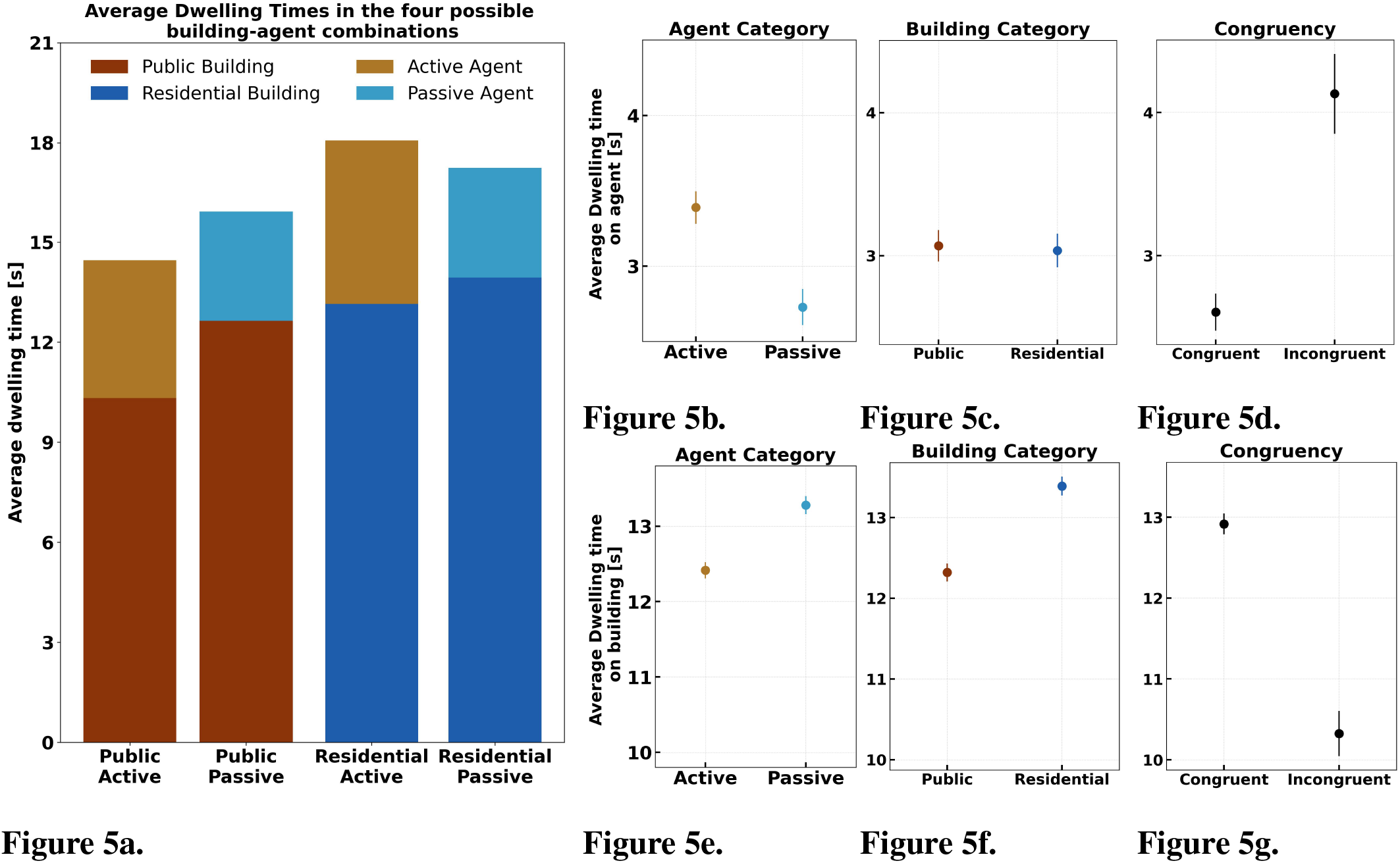
Dwelling times across Sessions: This figure presents the dwelling time, defined as the cumulative sum of fixations on specific objects during the exploration phase, measured in seconds. The bar graph shows the interplay of two factors, building and agent category. **(A)** illustrates the average dwelling times for four distinct building and agent type combinations. Each bar represents a unique combination: public buildings with active agents, public buildings with passive agents, residential buildings with active agents, and residential buildings with passive agents. The height of each bar indicates the average dwelling time, reflecting the relative visual attention each combination received. To the right, we examine one factor at a time for dwelling time on the agent on the top (panels **(B), (C)**, and **(D)**), and on buildings on the bottom (panels **(E), (F)**, and **(G)**).

Regarding building type, dwelling time on the agent was not significantly affected by the surrounding area, as agents in public areas (*M* = 3.07, *SD* = 3.07) and residential areas (*M* = 3.03, *SD* = 3.37, see Figure.5c) received similar attention. However, participants spent more time gazing at residential buildings (*M* = 13.39, *SD* = 9.01) compared to public buildings (*M* = 12.32, *SD* = 8.42, see Figure.5f). These results imply that incongruent agents divert focus from surroundings, as indicated by the anti-correlation in dwelling times between agents and buildings.

Examining congruency (see Fig. 5D), participants looked at incongruent agents (*M* = 3.23, *SD* = 3.48, see Figure.5d) for longer periods compared to congruent agents (*M* = 2.60, *SD* = 2.44). This pattern was opposite for building gazing time, with participants spending more time looking at buildings and surroundings when the agent matched the context in which it was placed (*M* = 12.92, *SD* = 7.65) compared to when the agent did not match the surroundings (*M* = 12.83, *SD* = 9.15, see Figure.5g). These results imply that when faced with incongruent agents, participants redirect their focus away from the surroundings, as evidenced by the anti-correlation in dwelling times between agents and buildings in the congruency factor.

To account for the high inter-individual variability and the nested structure of the data, we employed a linear mixed-effects model to predict dwelling time on agents, with subjects as a random effect. The fixed effects included the context (residential vs. public), agent type level (passive vs. active), the congruency of the agent with their surroundings (not congruent vs. congruent), and the interaction between agent type and context. The results indicated that participants gazed at agents for a significantly shorter period in residential contexts (*β*_Building_ = −0.41, *SE* = 0.08, *t* = −5.30, *p <* 0.001). Additionally, participants spent more time gazing at active agents compared to passive agents (*β*_Agent_ = 1.32, *SE* = 0.08, *t* = 16.94, *p <* 0.001). The congruency between the agent and its context also had a significant effect, with participants gazing at congruent agents for a shorter duration (*β*_Congruency_ = −0.58, *SE* = 0.13, *t* = −4.59, *p <* 0.001). Moreover, the interaction between context and agent action level was significant (*β* = −0.76, *SE* = 0.16, *t* = −4.88, *p <* 0.001), suggesting that the longest dwelling times were observed for active agents in residential settings. The findings reveal that residential active agents capture the most attention, especially when they clash with their surroundings.

We fitted an analogous linear mixed-effects model for the dwelling time on buildings. We found that participants gazed at buildings for a significantly shorter period in residential contexts (*β* = −2.02, *SE* = 0.25, *t* = −8.19, *p <* 0.001). Additionally, participants spent less time gazing at buildings with active agents compared to those with passive agents (*β* = −1.44, *SE* = 0.25, *t* = −5.83, *p <* 0.001). The congruency between the agent and the building also had a significant effect, with participants gazing at buildings with congruent agents for a longer duration (*β* = 3.07, *SE* = 0.39, *t* = 7.77, *p <* 0.001). Moreover, the interaction between context and agent action level was significant (*β* = −1.61, *SE* = 0.49, *t* = −3.26, *p* = 0.001), indicating that the shortest dwelling times were observed for buildings with active agents in residential settings. The data reveal that attention shifts to the agent when there’s incongruency with the context, but when congruency exists, attention is directed to the building.

### 2.2 Testing for spatial knowledge acquisition

We assessed participants’ spatial knowledge acquisition using pointing tasks. In the *Pointing to Buildings task*, participants were given screenshots of target buildings, while in the *Pointing to Agents task*, they received screenshots of agents against a grey background. Participants were asked to point toward the target. Accuracy was measured by calculating the angular difference between their pointing direction and the direction to the center of the target building or agent. This angular error served as the performance indicator, with a perfect score yielding zero degrees of error and higher values indicating greater inaccuracy.

#### 2.2.1 Pointing to Buildings

We hypothesized that participants would have more accurate pointing for public buildings, especially when targets had active, contextually congruent agent-building pairs. We applied a linear mixed-effects model to predict pointing errors, incorporating fixed and random effects. The random effects accounted for the test location of the pointing task (28 distinct locations with repeated measures). The fixed effects included the type of building, type of agent, congruency pairs, and the interaction between agent and building. The analysis revealed that pointing accuracy was significantly better, with lower errors in public buildings (*β* = −5.51, *SE* = 1.69, *t* = −3.28, *p <* .001). Active agents significantly improved pointing accuracy compared to passive agents (*β* = −7.17, *SE* = 1.72, *t* = −4.17, *p <* .001). The congruency of agent actions also played a crucial role, with incongruent pairs (where agent actions did not match the context) leading to better performance than congruent pairs (*β* = 6.88, *SE* = 1.97, *t* = 3.50, *p <* .001). Additionally, the interaction between agent and building type was non-significant, indicating that the main factor captured the relevant information regarding performance (*β* = 4.03, *SE* = 2.45, *t* = 1.64, *p* = .101). These results suggest that conditions where agents attract more visual attention also enhance pointing accuracy, particularly with active, contextually incongruent agents in public buildings. Thus, memorable human agents can boost spatial recall.

After the linear mixed-effects analysis, we examined the estimated marginal means (EMM) to clarify how different factors influenced pointing accuracy. Public buildings (EMM = 47.4, *SE* = 2.81) resulted in significantly lower errors than residential buildings (EMM = 50.9, *SE* = 2.94). Passive agents were associated with higher errors (EMM = 51.7, *SE* = 2.93) compared to active agents (EMM = 46.6, *SE* = 2.82). The interaction between agent and building type revealed that in residential contexts, passive agents resulted in the highest errors (EMM = 54.5, *SE* = 3.03), while active agents in residential contexts showed lower errors (EMM = 46.8, *SE* = 2.77). In public contexts, passive agents had higher errors (EMM = 49.2, *SE* = 2.75), while active agents in public buildings showed the lowest errors (EMM = 45.8, *SE* = 2.81). The congruency of agent actions also played a key role, with incongruent pairs performing better than congruent pairs. Specifically, incongruent pairs had lower errors (EMM = 45.7, *SE* = 2.81) compared to congruent pairs (EMM = 52.6, *SE* = 3.14). The findings emphasize that active, contextually incongruent agents improve pointing accuracy, reducing errors and equalizing performance across building types (see Figure.6a).

**Figure 6.**
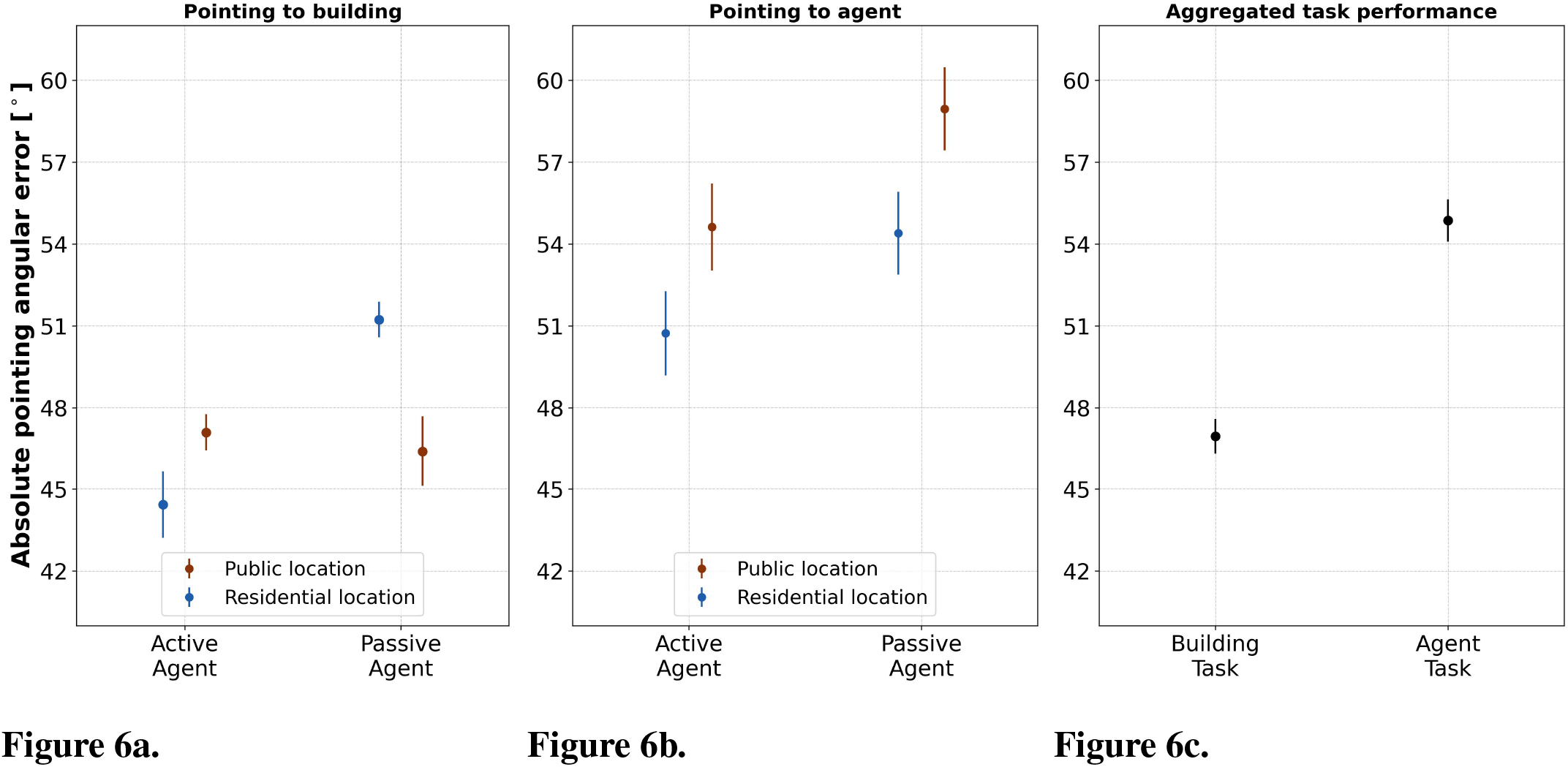
Fig. 6: Analysis of Pointing Accuracy across experimental conditions. This figure illustrates the absolute pointing angular error, a measure of spatial recall accuracy whose lower values indicate better performance, with data points representing the mean error rates and error bars indicating standard errors. **(A)** Pointing to Buildings: This plot delineates participants’ pointing accuracy to buildings paired with active and passive agents. **(B)** Pointing to Agents: Here, participants’ accuracy in pointing to agents is compared, with a focus on the agent’s activity level and the type of building. **(C)** Aggregated Task Performance: This plot displays overall performance across both building and agent pointing tasks.

#### 2.2.2 Pointing to Agents

We hypothesized that participants would demonstrate better pointing accuracy for the locations of active agents, particularly those positioned in front of public buildings. To test this hypothesis, we employed a linear mixed-effects model similar to the one used for analyzing pointing-to-building performance. The model included crossed random effects for subjects and the starting locations of the pointing tasks, covering 28 distinct locations. The analysis revealed significant main effects for both the building type (*β* = −4.76, *SE* = 1.50, *t* = −3.17, *p* = .002) and agent category (*β* = 4.22, *SE* = 1.51, *t* = 2.80, *p* = .005). However, the interaction between context and agent action was non-significant (*β* = −0.79, *SE* = 3.01, *t* = −0.26, *p* = .792).

Contrary to our hypothesis, participants were more accurate in pointing to agents located at residential buildings (EMM = 52.1, *SE* = 3.22) rather than public ones (EMM = 56.3, *SE* = 3.21), despite being more accurate overall when pointing at active agents (EMM = 51.8, *SE* = 3.23) compared to passive agents (EMM = 56.6, *SE* = 3.21, see Figure.6b for the observed averages). The findings indicate that task accuracy is differentially influenced by the recall of agents and their surroundings. Specifically, reduced visual attention on agents hinders their spatial recall, while for buildings, participants benefit from attention on both the agent and the building.

#### 2.2.3 Accuracy differences between pointing to buildings and pointing to agents

We expected participants to be less precise when pointing to agent stimuli compared to building stimuli. To investigate this, we first assessed whether participants exhibited significantly lower precision when pointing to agents. This was achieved by fitting a two-crossed random effects model (i.e., ID and pointing location) and predicting pointing error with the type of stimuli (agent vs. building) as the sole predictor. The analysis revealed that participants were indeed significantly less precise when pointing to agent stimuli (*β* = −7.74, *SE* = 0.98, *p <* .001) compared to their performance in the pointing-to-building task. The estimated marginal means (EMMeans) showed that participants had higher pointing errors for agent stimuli (EMM = 54.4, *SE* = 2.55) compared to building stimuli (EMM = 46.7, *SE* = 2.53). This clear difference is illustrated in Figure.6c, where the average performance indicates that participants could recall the precise locations of buildings more accurately than those of human agents. The data reveal that while building locations were recalled more accurately, agents may have served as a salient feature, enhancing the overall spatial recall even if their precise positions were less accurately remembered.

#### 2.2.4 Inclusion of gaze as a predictor of performance

We hypothesized that the time spent looking at both agents and buildings would significantly predict participants’ performance, as both could serve as recall cues for location. To assess this hypothesis, we incorporated the dwelling time in seconds each participant spent gazing at the agents and buildings as fixed effects in the pointing-to-building and pointing-to-agent tasks (see Figure.7). For the pointing-to-building task, the results showed that only the dwelling time on buildings significantly predicted performance (*β* = −0.18, *SE* = 0.05, *t* = −3.62, *p <* .001), while the dwelling time on agents did not have a significant effect (*β* = −0.27, *SE* = 0.15, *t* = −1.79, *p* = .073). The type of agent (active vs. passive) remained a significant predictor, with active agents leading to better performance (*β* = −6.79, *SE* = 1.74, *t* = −3.91, *p <* .001). The type of building also remained significant (*β* = −5.71, *SE* = 1.68, *t* = −3.41, *p <* .001). Additionally, the congruency between the agent’s actions and the building showed a significant effect (*β* = 7.24, *SE* = 1.98, *t* = 3.66, *p <* .001). In the pointing-to-agent task, the results revealed that neither the dwelling time on agents (*β* = −0.41, *SE* = 0.24, *t* = −1.72, *p* = .086) nor the dwelling time on buildings (*β* = −0.002, *SE* = 0.09, *t* = −0.02, *p* = .981) were significant predictors of performance. However, building type, with the opposite pattern as in pointing to buildings (i.e., residential locations being better remembered, *β* = 3.97, *SE* = 1.53, *t* = 2.60, *p* = .009) and agent type, with the same pattern (i.e., active agents being better remembered, *β* = −4.20, *SE* = 1.54, *t* = −2.73, *p* = .006) remained significant, while their interaction was not (*β* = −1.25, *SE* = 3.03, *t* = −0.41, *p* = .680).

**Figure 7.**
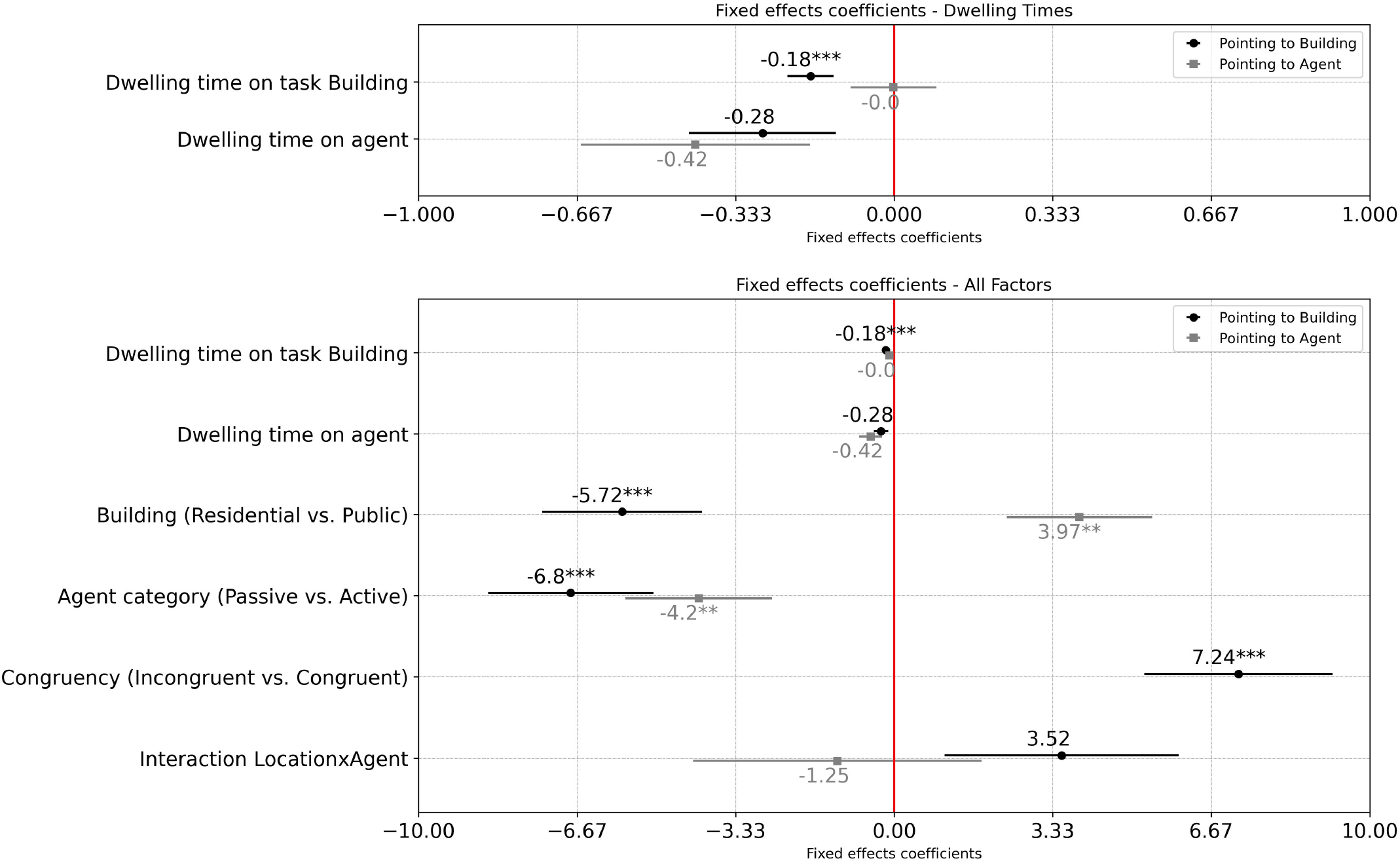
Accuracy prediction models: The estimates for the five effects and one interaction are displayed using one standard error as error bars. The linear mixed models were computed employing a structure of two crossed-random effects, which accounted for the pointing place (i.e., the 28 pointing locations) and the incorporation of repeated measures. The red line denotes no difference amongst the levels of the factor. Instances marked by three asterisks indicate significance levels falling below .001, while two asterisks indicate significance below .01. The upper panel represents the same results as the first two lines of the lower panel but zoomed in.

**Figure 8.**
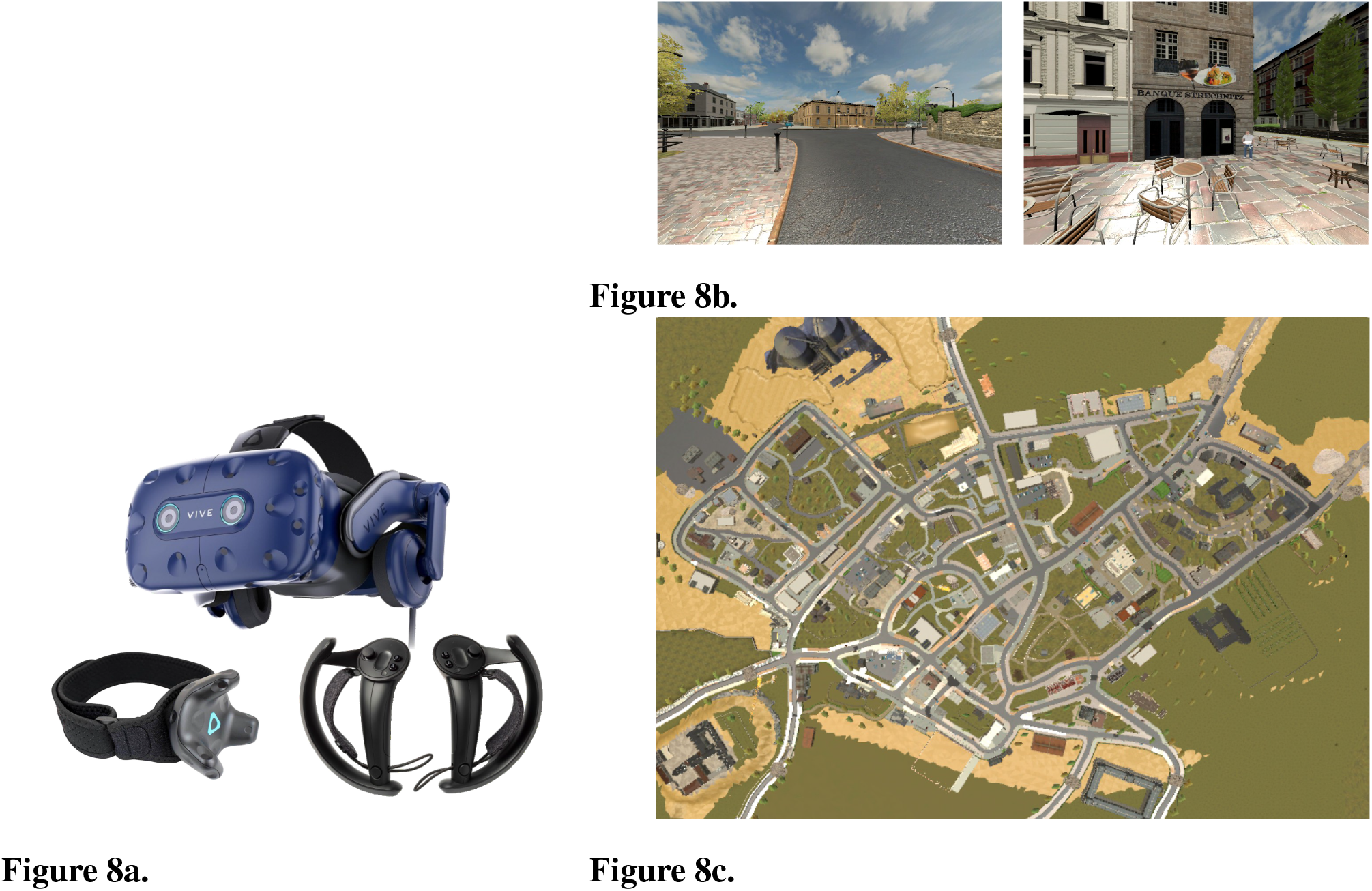
Overview of the Virtual Reality Environment. **(A)** Experimental setup, the equipment was used with a swivel chair for lateral navigation and a joystick on the VIVE controller for forward movement. **(B)** First-person immersive view available to participants. **(C)** Aerial view of the virtual city.

Comparing the results from the two tasks, it becomes evident that there is an inverse relationship between the ability to locate agents and buildings, as both compete for attention. Active agents improved the recall of their locations, yet the presence of public buildings appeared to detract from the ability to remember the agent. Therefore, while agents may serve as useful proxies for recalling target locations, their own positions are not necessarily better remembered in relation to their surroundings.

## 3 DISCUSSION

This study explored the impact of human agents on spatial navigation and knowledge acquisition within a controlled virtual reality city environment. Our findings suggest that while the presence of agents influenced spatial exploration, this effect was relatively minor, as the overall exploration strategy remained largely unchanged. Nonetheless, agents increased the likelihood of participants revisiting the buildings where they were located upon first exploration of the environment. In these spaces, participants paid more visual attention to agents that did not match their surroundings, indicating a context-dependent perception of these agents. During exploration, agents and buildings competed for attention, shaping how spatial information was processed. When testing spatial knowledge, pointing accuracy for buildings benefited from both the buildings and the agents, suggesting a synergistic effect where agents enhanced the salience of building locations. However, this benefit did not extend to pointing accuracy for agents, where accuracy did not improve when the agent was located at a more salient building. Additionally, contextual incongruence played a crucial role in enhancing pointing accuracy, highlighting non-linear effects that further underscore the complexity of how the presence of others can shape salience. Overall, it can be concluded that the presence of agents and their integration into the environment significantly shape how spatial information is remembered and, to a lesser extent, how the space is explored.

### 3.1 Limitations

A key limitation of this study is that the agents, including those classified as “active,” remained static, which does not reflect the dynamic interactions typical of real-world scenarios. Additionally, the fixed positions of the agents throughout the study may have obscured more subtle changes in participant behavior that could occur with moving or varying agent locations. This lack of movement may limit the generalizability of our findings to real-life situations where the presence and behavior of others are more variable. However, we maintained this static feature to ensure consistency between the two agent categories, allowing us to investigate how even minimal actions, such as holding an object, can influence participant behavior and performance. Through this design, we identified a minimal level of experimental manipulation that exerts a differential impact by human agents without introducing the inherent eye movement biases that a moving group might create. It is necessary to note that people’s interactions with agents can differ based on the agent’s group size, and our study focused solely on single agents. We excluded groups because previous research shows that individuals are more inclined to approach single agents Bonsch et al., 2018, and crowded spaces may deter pedestrians, causing navigation avoidance Dickinson et al., 2019; Li et al., 2019. Our experiment, with 56 agents per square kilometer, likely stayed below the discomfort threshold. Therefore, our findings suggest that lower agent densities can be utilized to study navigation behaviors without inducing avoidance, providing insights into optimal crowd levels for effective spatial navigation.

The virtual environment, though extensive, cannot fully replicate the complexity and nuances of real-world navigation and interactions, particularly concerning general movement flow. In real-world environments, a multitude of factors contribute to how individuals navigate and perceive spaces, including moving objects, varying light conditions, and the presence of other moving entities. While controlled and consistent, the static nature of our virtual environment lacks these dynamic elements that typically influence human behavior. This limitation means that our study did not capture certain interactive and responsive behaviors, which are naturally triggered by a dynamic environment. However, we deliberately opted against including dynamic environmental elements for two primary reasons: first, to optimize the sample rate, as additional movement could burden the system and reduce the reliability of data capture; second, to avoid introducing confounding variables that might draw visual attention and skew our eye-tracking analysis.

### 3.2 Agent impact on spatial exploration

To understand participants’ broader navigational behavior, we analyzed their spatial coverage and decision-making strategies, contrasting participants who explored the city with (our experiments) and without agents (control group from (Schmidt et al., 2023)). We developed a behaviorally-based method to quantify the exploration trade-off through a primal graph that captures decision-level spatial navigation dynamics in the virtual city environment. The results demonstrated that as participants became more familiar with the environment, they progressively explored more of the city with each session. Notably, participants showed consistent patterns in both the total area covered and the balance between exploratory and conservative behavior, regardless of whether agents were present. Our findings align with previous research on exploration-exploitation dynamics. For instance, Choi et al. (2012) observed that participants switched from long, unpredictable search movements (Levy flights) when uncertain to shorter, deliberate paths (Brownian walks) as they gained confidence. Similarly, (Dickinson et al., 2019; Li et al., 2019) demonstrated that various optimal foraging strategies, from ballistic to Brownian motion, emerge based on the experience with the environment. In concordance with these free exploration studies, our study found that participants’ strategies evolved in response to their growing familiarity with the VR city, balancing exploration and exploitation with agents not interfering in this process. By analyzing participants’ navigational decisions, specifically their tendencies between revisiting known locations and exploring new paths, we identified an enhanced likelihood of revisiting behavior unique to the first session, where participants were significantly more likely to move in the direction of agents. These findings suggest that while agents may have an initial biasing effect on participants, they do not substantially influence broader spatial exploration strategies. Globally, participants maintained a consistent exploration rate, characterized by a mix of exploratory and conservative decisions, across all sessions. This consistency reinforces the idea that agents exert a localized impact on navigation, affecting behavior at specific decision points without altering the overall approach to exploring the environment.

### 3.3 Visual exploration of the environment

When participants explored areas with agents, they preferred visually engaging with active agents over passive ones despite all agents being static. This aligns with theories indicating that stimuli with higher informational density demand more cognitive processing, making them more salient Henderson (2007); Summerfield and Egner (2009); Wolfe (2020). Such stimuli are perceived as more salient particularly when they contain elements relevant to the task at hand or the environment’s narrative (Harel et al., 2014; Miller et al., 2014). Previous studies have found that, whether consciously or not, people map their surroundings in terms of future action planning (Bach et al., 2014; Bonner and Epstein, 2017; Ohm et al., 2014). This tendency extends to situations involving other people, where the presence of a person in a position to interact with objects can lead observers to adopt that person’s spatial perspective (Tversky and Hard, 2009). Thus, one could argue that active agents holding objects and displaying varied body positions garner longer dwelling times because they hint at navigational affordances; that is, they prime the participants for possible actionable routes. Consequently, active agents would have higher informational value for participants navigating a city since they hint at contextual elements that could be acted upon.

Furthermore, our analysis revealed that the longest gaze times were directed toward active agents that mismatched their surroundings. This finding can be understood through the framework of spatial schema, which are structured bodies of prior knowledge used to navigate and interpret environments (Farzanfar et al., 2023). When these agents clashed with their background, it likely elicited an expectancy violation (Burgoon, 2015), prompting participants to engage in a longer visual search to reconcile this disparity. Together, this supports the notion that agents are perceived contextually, and participants showed signs of trying to integrate them into the space they inhabit.

Regarding building gazing, our participants behaved contrary to our initial hypothesis, spending more time inspecting residential instead of public buildings. This finding contradicts previous research, which generally suggests that people tend to spend longer looking at public buildings like restaurants, stores, and landmarks due to their visual complexity and social relevance (Dalton et al., 2019; Rounds et al., 2020). Eye-tracking research has consistently found that functional and visually salient landmarks, such as public buildings, are more likely to be used for navigation and remembered longer Farran et al., 2016; Walter et al., 2022. However, our participants’ preference for residential buildings suggests a potential contextual influence. In our study, all task-relevant buildings had graffiti, which may have caused the extended gaze time. Additionally, all task-relevant residential buildings had an agent in front of them, which could have garnered more viewing time. Finally, it might be the case that since the residential buildings were more similar to each other, participants required more time to tell them apart. While one gaze at a donut shop can already provide a clear memory cue, a gaze at a standard residential house might need more time to gather a cue that would help differentiate it from neighboring buildings.

### 3.4 Effect of Agents on Spatial Knowledge Acquisition

Our study provides significant insights into the role of human agents in spatial knowledge acquisition, particularly regarding pointing accuracy. The consistent finding across both pointing-to-agent and pointing-to-building tasks is that active agents significantly enhance performance. This aligns with previous research indicating that social targets and interactive cues improve navigational accuracy and spatial encoding. For instance, Kuehn et al. (2018) demonstrated that participants exhibited reduced positional errors when navigating with a person as a target, suggesting that social targets facilitate spatial encoding by enhancing the processing of both body-based and environment-based cues. Similarly, Gunalp et al. (2019)) found that including an avatar in spatial perspective-taking tasks improved performance compared to abstract directional cues, underscoring the importance of social and interactive aspects in aiding mental simulation processes required for spatial tasks. More specifically, the pronounced effect of active agents on spatial knowledge acquisition may be attributed not only to increased visual engagement but also to the cognitive implications of perceived action. Previous studies have shown that even the suggestion of action in still images can enhance spatial processing (Tversky and Hard, 2009). In our study, when those actions were incongruent with the surroundings, performance was further enhanced, likely because reconciling the mismatch strengthened the memory cues for the location. This suggests that by implying potential actions, active agents encourage participants to process spatial information from multiple perspectives, thereby enhancing their overall spatial understanding and memory.

A notable finding from our study was the overall higher precision in pointing to buildings compared to agents. This indicates that while human agents can enhance spatial recall, participants generally exhibited greater accuracy when recalling traditionally static buildings. This could be attributed to buildings’ more stable and distinctive features, which provide consistent spatial cues. In contrast, agents, being potential sources of movement, may introduce variability in spatial memory. The differential impact of agents on pointing accuracy to buildings and agents highlights the importance of contextual relevance in spatial knowledge acquisition. Active agents, particularly those performing contextually incongruent actions, appear to be strong spatial anchors for their surroundings but are not remembered as landmarks.

### 3.5 Implications for Spatial Navigation Systems

These findings have significant implications for the design of spatial navigation systems. Incorporating human contextually relevant cues such as active agents can enhance spatial recall and bias navigation efficiency. This aligns with the broader literature on wayfinding, which emphasizes the importance of functional landmarks and visual salience (Franke and Schweikart, 2017; Ohm et al., 2014). By leveraging active agents and ensuring their actions are contextually relevant, navigation systems can provide more effective spatial cues, aiding users in more accurately recalling and navigating environments. Additionally, intentionally contrasting agents with the environment could improve the memory of a location as it might contradict expectations. In summary, our study underscores the critical role of human agents in shaping spatial knowledge acquisition. Active agents enhance pointing accuracy and spatial recall, mainly when they are contextually relevant. These findings contribute to a deeper understanding of how human elements within an environment can be effective navigational aids, providing valuable insights for developing more intuitive and effective spatial navigation systems.

### 3.6 Conclusion

This study explored the impact of human agents on spatial navigation and knowledge acquisition within a virtual city. Our findings show evidence that human agents, particularly those portraying actions, locally influence navigational behavior and enhance spatial recall. Active agents drew more visual attention and served as effective spatial cues, improving participants’ ability to recall specific locations. Interestingly, agents that were incongruent with their surroundings further enhanced spatial memory, suggesting that expectancy violations prompt spatial knowledge acquisition. These results underscore the importance of incorporating human elements into virtual environments to better understand their role in spatial cognition. The insights gained from this study have potential applications in designing more intuitive spatial navigation systems and enhancing training programs for environments where human interaction is a critical component.

## 4 METHODS

### 4.1 Participants

We recruited 70 participants, distributing them equally across two experiments (35 per experiment). Participants were required to have normal or corrected-to-normal vision and to attend five 30-minute sessions, plus one final 60-minute test session in a virtual reality (VR) environment. Sessions were scheduled at intervals ranging from four hours to three days. However, attrition occurred due to sickness or missed appointments, resulting in 10 participants being unable to complete the required schedule and 15 participants being excluded due to technical failures, like interruption of files. Additionally, three participants withdrew due to motion sickness. The final sample included 21 participants (12 females, *M*_age_ = 25.33 years, *SD*_age_ = 7.66) for the first experiment and 21 participants (11 females, *M*_age_ = 22.31 years, *SD*_age_ = 2.80) for the second experiment. All participants provided written informed consent. Compensation was given in the form of “participant hours,” a common requirement within the study programs at the University of Osnabrü ck, where the ethics committee approved the study following the ethical standards of the Institutional and National Research Committees.

### 4.2 VR environment: Westbrook

The virtual environment, known as Westbrook, was developed using Unity LTS 2019.4.27f1, as described by Schmidt et al. (2023). The virtual city covers an area of approximately one km^2^ and is inspired by the layout of the Swiss city Baulmes. It features 236 buildings, including 26 public buildings (i.e. shops, basketball courts), 26 residential buildings marked with graffiti, and 180 regular buildings devoid of graffiti. Additionally, four large buildings are strategically placed at the city’s periphery. The buildings with graffiti were designated as task buildings in both our experiment and the experiment by Schmidt et al. (2023) and are distributed roughly equally throughout the city. Mesh colliders were applied to all objects to facilitate the tracking of participants’ visual behavior through 3D eye vector projection onto the environment.

Navigation within the city is confined to visible paths and streets defined by a custom navigation mesh, with restricted access to areas blocked by fences or other physical barriers. Conventional orientation cues such as street names, house numbers, and solar positioning were deliberately excluded to enhance the navigational challenge. Participants controlled their translational movement up to a maximum speed of 5 m/h using a joystick, while rotational movements were facilitated by physical rotation on a swivel chair.

### 4.3 The Human Agents

The inclusion of 56 human agents from the Adobe Mixamo collection Mixamo, 2008 was designed to explore the impact of human elements on spatial learning. Agents were divided into two groups: 28 (14 males and 14 females) in the active agent group and an equal number in the passive agent group. Active agents were equipped with items and body postures that underscored the significance of specific buildings—such as a sandwich at a sandwich shop or a toolbox at a tool store (see Figure.9a). Conversely, passive agents, matched in skin tone, hair color, and gender with the active group, were depicted in a relaxed standing pose see Figure.9b), enhancing environmental realism without engaging in explicit, building-specific activities. All agents were static and non-responsive to the participants.

**Figure 9.**
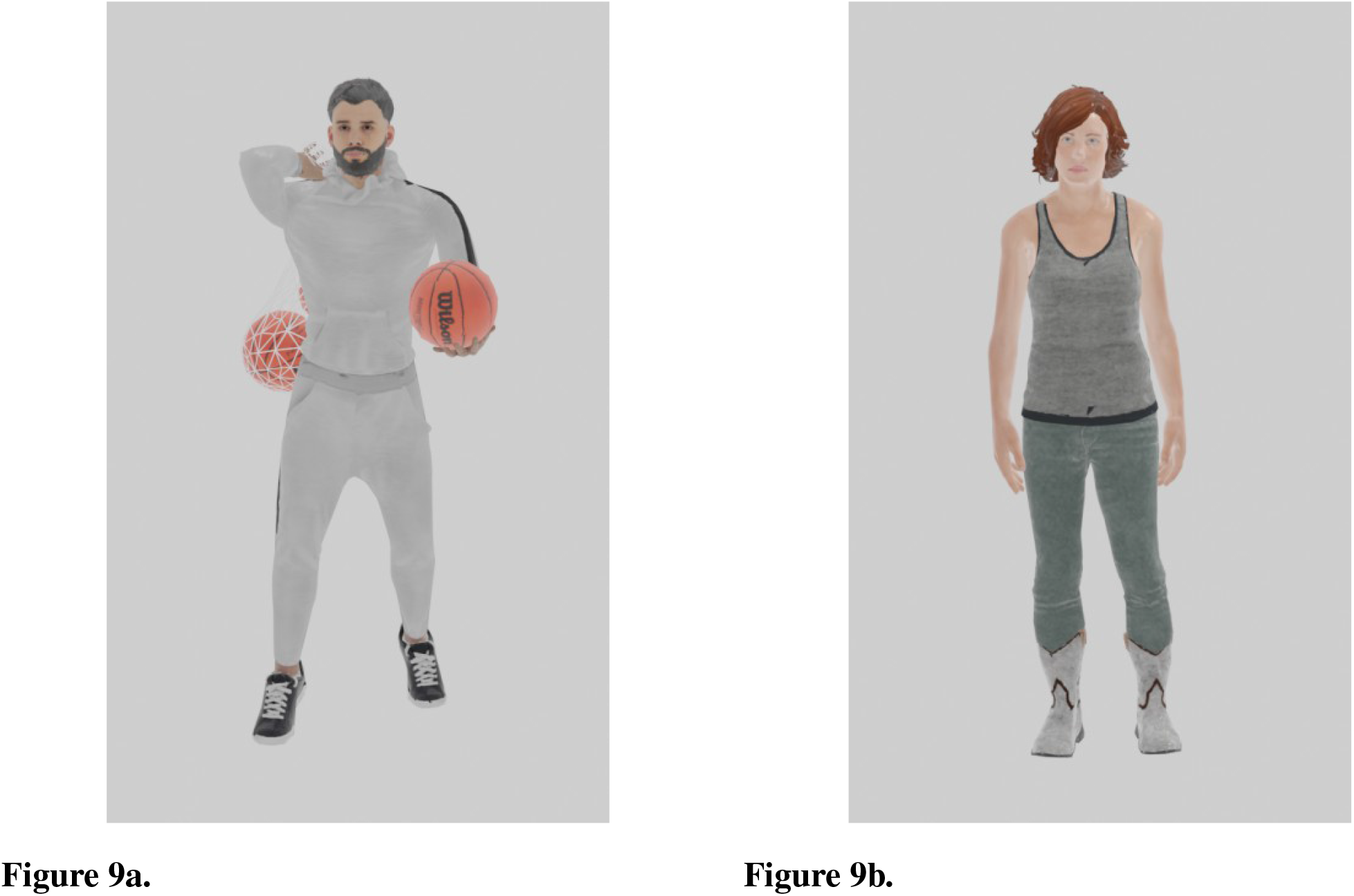
Human Agents. **(A)** Active human agent. **(B)** Passive human agent.

In the first experiment, active agents were designed to be positioned in public buildings to reflect the thematic context of their surroundings. Specifically, active agents were positioned in front of the 26 street art-marked public buildings and two larger buildings. For instance, a construction worker wielding a shovel was situated at a construction site among 26 public buildings, along with two thematic global landmarks (i.e., a church-goer with a bible at a church and another construction worker at a castle under construction). Conversely, passive agents were stationed in neutral stances within the 26 residential areas and at two additional larger buildings (i.e., a silo and a windmill).

In the second experiment, we changed the setup by redistributing active and passive agents across public and residential areas, intentionally breaking the congruency established in the first experiment. This shuffle placed half of the active agents in non-matching public buildings and the other half in residential settings. Similarly, passive agents were also re positioned, with half now found in public spaces and the remaining half in residential areas. This reconfiguration aimed to probe the effects of agent-building incongruency on spatial learning, challenging the direct association between agents and specific buildings.

### 4.4 Experimental procedure

Both experiments were divided into two main phases: the exploration and task assessment phases. The exploration phase consisted of five sessions, each lasting 30 minutes, resulting in 150 minutes of free exploration (see Figure.1). The task assessment phase was the sixth session, during which participants performed a pointing task. Sessions were spaced with intervals ranging from a minimum of four hours to a maximum of three days.

In the initial session, participants received a comprehensive overview of the experiment and provided written informed consent. They then completed the “Fragebogen Räumlicher Strategien” (FRS) questionnaire to assess their spatial orientation abilities (Münzer and Hölscher, 2011). Subsequently, participants were familiarized with the VR environment in a neutral VR room. In this room, they practiced controlling their lateral movements by rotating in a swivel chair and regulating their forward movement using the joystick on their VIVE controller. The exploration phase within the actual city began once participants confirmed their understanding of the movement mechanics.

During the exploration phase, participants were instructed to freely explore the virtual city for 30 minutes, imagining they were getting acquainted with a real city they would be tested on by the end of the experiment. In contrast with Schmidt et al. (2023), we did not instruct them to look for street art specifically marked houses, but rather to just try to pay attention to the city’s layout. The sessions were divided into three 10-minute segments, and a 9-point eye-tracking calibration and validation were performed at the start of each segment to ensure measurement accuracy, maintaining a visual tracking error of less than one degree.

Participants performed a pointing task within the VR environment in the assessment session from a first-person perspective. Within the task, participants were teleported to 28 predetermined locations inside Westbrook. At each location, they were presented with a target at the center of their visual field and instructed to point directly at the center of the target. In experiment 1, the targets were solely buildings (i.e., pointing-to-building). In contrast, in experiment 2, we included a test that displayed only the agent against a gray background (i.e., pointing to agents). To account for the sequence effect, the task order was strategically randomized, with one-half of the participants starting with the pointing-to-agent task followed by the pointing-to-building task and the other completing the tasks in reverse order. Finally, all participants completed three self-assessment questionnaires that inquired about their perception of each agent, their social tendencies in real life, and the realism level of the VR. The questionnaire data has not been analyzed in this work.

### 4.5 Experimental setup

The exploration and task assessment sessions were conducted with a desktop computer with an Intel(R) Xeon(R) W-2133 CPU, 16 GB RAM, and a Nvidia RTX 2080 Ti graphics card. The VR environment was rendered with a HTC Vive Pro Eye head-mounted display (HMD) setup with a refresh rate of 90Hz and horizontal and vertical field of view of 106° and 110°, respectively. We used four SteamVR Base Stations 2.0, an HTC VIVE body tracker 2.0, and Valve Index controllers to monitor participants’ positions within the environment. This combined setup achieved sub-millimeter precision in capturing the head, body, and eye positions, as well as rotation and orientation.

### 4.6 Spatial task

Participants performed pointing tasks in VR from a first-person perspective. In the pointing-to-building task, they were teleported to 28 unique locations within the city, each serving as a distinct reference point. The sequence of these locations was randomized for each participant to prevent order effects. At each location, participants pointed repeatedly toward one of 56 potential targets, represented by static images of buildings, with agents positioned in front of them. These images were captured perpendicularly at a height of 1.80 meters to ensure consistency in visual presentation.

The trials began with a visual and auditory cue: a 25ms green circular loading bar at the screen’s center accompanied by a beep. The target image appeared at the upper center of the screen, with a green dashed laser beam providing a visual guide for pointing. Participants indicated their direction by pressing a trigger button on their controllers. Each trial was timed for 30 seconds, automatically concluding if a direction was not indicated within this period. Performance was assessed by measuring the angular difference between the participant’s pointing direction and the precise center vector of the target location.

In the initial experiment, participants completed 336 trials, pointing at 12 unique targets from each of the 28 reference locations. The reduction in the number of trials in the second experiment, where participants completed only 224 trials, pointing at eight distinct targets from each location, was necessitated by the additional pointing to agents task. In both experiments, the trials were balanced to ensure an even distribution of target types: 50% directed participants to public locations and 50% to residential areas.

### 4.7 Pointing to Agents Task

This task was fundamentally equivalent to the “Pointing to Buildings Task” but specifically focused on agents as the target. The targets featured centered screenshots of individual agents set against a grey backdrop, captured perpendicularly at a height of 1.80 meters. Participants undertook 224 trials and were instructed to point at eight unique agent targets from each of the 28 reference locations. The task mirrored the structure of the building task in terms of target placement, visual and auditory cues, and time constraints to maintain consistency in the testing conditions. Additionally, the sequence of these pointing locations was randomized for each participant, with the goal of avoiding order effects and preserving the integrity of the experimental data.

### 4.8 “Fragebogen Rä umlicher Strategien” (FRS) questionnaire

The “Fragebogen Räumlicher Strategien” (FRS; Mü nzer and Hö lscher (2011)) questionnaire is a 7-point Likert scale that asks the participants to estimate their spatial orientation abilities in three areas of spatial knowledge in real-world scenarios. First, the global sub-scale consisted of 10 items (*α* = 0.89) inquiring about the subject’s ability to navigate routes from an egocentric perspective. The survey sub-scale incorporates seven items (*α* = 0.87) focused on the subject’s ability for mental mapping from an allocentric perspective. The cardinal sub-scale comprises two items (*α* = 0.80) that query the ability to point toward cardinal points. The FRS measures participants’ likelihood to apply spatial strategies related to egocentric/global knowledge, survey knowledge, or cardinal directions, respectively.

### 4.9 Movement tracking in the city

#### 4.9.1 Navigational tracking

To analyze participant trajectories and their exploration decisions, we created a data-driven graph based on their actual trajectories while exploring the virtual city. Thus, this graph reflects only the paths and areas that participants walked through, constituted by the streets (edges) and crossings or decision points (nodes) that participants walked through. We first generated a heatmap of participants’ movement to transition from raw sequential coordinates to spatial data. This heatmap accounted for the number of times a participant stood at a specific cell within a defined 4m × 4m grid on top of the city map. This heatmap was then turned into a binary image, where cells visited at least once were assigned a value of one and cells without visits a value of zero. The resulting image clearly outlined the walkable paths and connections within the city. We filled isolated holes within the streets to ensure the algorithm generating the graph did not create extraneous nodes in these areas. We then generated a skeleton from the binary image, reducing the city’s representation to a one-pixel width while preserving its topography and connectivity. This skeleton was the foundation for identifying the graph’s nodes and edges. Using the external Python library “sknw”, we generated a NetworkX graph from the city skeleton image. Nodes were numbered sequentially, and edges were named based on the nodes they connected (see Figure.3a). For example, edge [76,60] connected nodes 76 and 60. Some manual adjustments, such as adding an edge, were necessary to perfect the graph. Each pixel in the skeleton was recognized as belonging to a node or an edge.,. This information was later used to plot the graph and calculate distances to identify the graph elements visited by participants during their sessions.

To analyze exploration strategies, we developed an algorithm that converts participants’ coordinates data into a sequence of visited nodes and edges. This process involved mapping the coordinates to the skeleton heatmap’s 4mx4m cells and determining the closest graph element using the Euclidean distance formula. The algorithm tracked transitions between edges and nodes, recording each visited element. This record included the time of entry, graph element type (node/edge), and number of visits to the graph element. As well as the available paths from the node of origin and their respective number of previous visits to the possible elements available from that position. We adjusted the nodes’ radii to match the width of the corresponding streets. This adjustment ensured accurate detection of participant presence at nodes, preventing unnoticed transitions between edges. Overlapping node radii in areas with several short streets were resolved by assigning participant positions to the nearest node centroid.

#### 4.9.2 Navigational Pattern Classification: Strategy Matrix

We condensed participants’ decisions at nodes into a strategy matrix to analyze their exploration patterns. The strategy matrix was organized with the number of visits to the chosen node on the rows and the number of visits to the not-chosen nodes on the columns. Each decision was recorded in the row corresponding to the number of previous visits to the selected path. We added one to each column corresponding to the number of prior visits on the other available paths from that decision point that were not selected. For instance, if a participant was at a node that had three direct neighbors, visited zero, five, and two times, and chose to move in the direction of the node visited two times, we would add a count of one to the positions [0,2] and [5,2]. This represents that the participant moved to a node with two visits over the options that had been visited zero and five times (see Figure.3b). Decisions above the diagonal line in the strategy matrix were considered conservative, indicating that participants preferred nodes they had visited more frequently in the previous example, the one added at [0,2].In contrast, decisions below the diagonal line were considered exploratory, as participants chose nodes with fewer visits compared to their neighboring nodes, in the example above the one added at [5,2]. Decisions exactly on the diagonal were neutral, as both the chosen and the adjacent nodes had the same number of previous visits. This method allowed us to quantify and compare participants’ exploratory and conservative tendencies as they navigated the virtual city, providing insight into their spatial decision-making processes as they gained experience inside Westbrook.

#### 4.9.3 Eye-tracking preprocessing and classification

We applied a velocity-based algorithm that classified continuous eye movements into gazes and saccades and corrected the resulting gazes for the participants’ movement in the 3D environment, as developed by (Nolte et al., 2024). In preparation for this algorithm, we preprocessed the data by excluding portions detected as invalid (e.g., blinks). In cases where more than one collider hit was detected within the same sample, we retained the closest hit to the participant, except for background colliders, such as leaves or fences, in which case we kept the second closest hit. As a last step, we dropped duplicated samples and applied a 5-point median filter to the gaze coordinates. The algorithm calculates the median velocity of eye movements within a time window (in our case, a 10-second window). Samples exceeding this velocity threshold are identified as saccades, while those falling below the threshold are classified as gazes. To ensure accuracy, we applied outlier detection based on median absolute deviation to correct for gaze events with anomalous durations (i.e., exceeding three median absolute deviations). Dwelling time was defined as the cumulative time a subject spent gazing at a specific object within the city across all sessions. We computed the dwelling time for each subject-object pair during their entire exploration period within the Westbrook environment.

### 4.10 Data Analysis

Given the hierarchical structure of our data, we employed Linear Mixed-Effects Models (LMMs) for our analysis. The modeling was done using R 4.3.2 with the lmer() function from the lme4 package. We used Restricted Maximum Likelihood (REML) for estimation and the nloptwrap optimizer (Bates et al., 2015). This approach allows us to handle the nested structure of our data, with random effects to account for within-subject variability. Fixed effects with two levels were effect-coded to ensure the betas reflect the differences between these levels, using the first level of each pair as the base for comparison.

#### 4.10.1 Exploration Phase Analysis: Navigational Coverage of the City

To analyze participants’ free exploration patterns, we quantified their walking behavior through the virtual city using a primal city graph Neal (2013). The coverage ratio, defined as the proportion of unique nodes visited during each session, was modeled using a linear mixed-effects approach. The model included fixed effects for the session, experiment, and interactions, with planned contrast testing for each session against the first. Random intercepts for participants were included to account for repeated measures within subjects. The model formula for the individual session coverage ratio was:

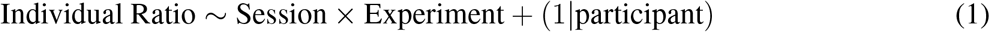

The same structure was used to test for the cumulative ratio of visited nodes, in which we kept track of how many unique decision points each participant had visited, accumulating them between sessions. The model formula for the cumulative coverage ratio was:

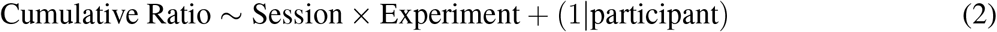

#### 4.10.2 Walking Strategies: Exploratory vs. Conservative Navigational Behavior

To investigate the extent to which participants used conservative or exploratory walking strategies, we modeled the number of decisions at each node. A linear mixed-effects model was used with the session, strategy (conservative vs. exploratory), and experiment as fixed effects and random intercepts for participants.

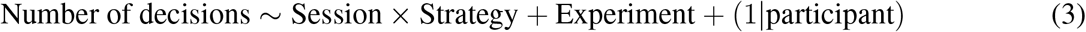

#### 4.10.3 Visual Behavior During Exploration

We quantified participants’ visual behavior by summing the cumulative time spent gazing at objects (dwelling time) during the entire exploration phase. Separate models predicted dwelling time on agents and buildings, with fixed effects for agent action level (passive vs. active), context effect (residential vs. public), the congruency of the agent with their surroundings (not congruent vs. congruent), and their interactions, including random effects for participants with the following formulas:

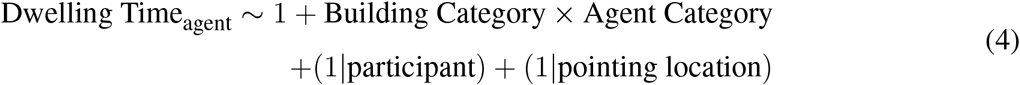

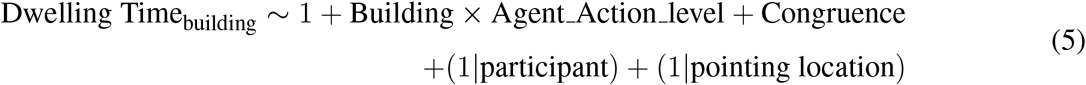

#### 4.10.4 Pointing Task: Pointing to Buildings

We assessed spatial knowledge using pointing tasks, calculating the angular error as the performance indicator. Two separate linear mixed-effects models predicted pointing error based on building type (residential vs. public), agent type (passive vs. active), the congruency of the agents with their surroundings, dwelling time on agents, dwelling time on buildings, and the interaction of agent and building type. Random effects for participants and pointing locations were included in each model:

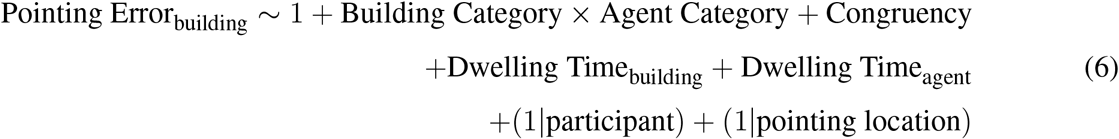

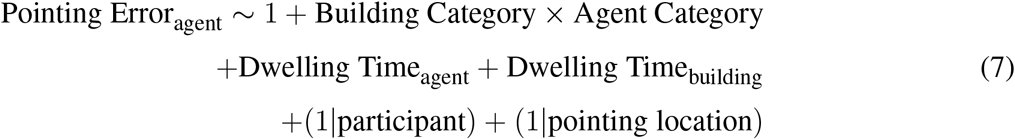

#### 4.10.5 Comparing Pointing Accuracy Between Tasks

To compare accuracy between pointing to buildings and pointing to agents, we used a model with the type of stimuli (agent vs. building) as the fixed effect and participants and pointing locations as random effects:

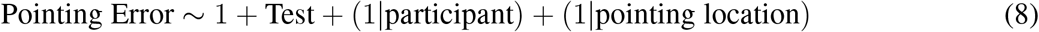

Following model fitting, we performed likelihood ratio tests to compare each model against a null model containing only the intercept, evaluating the added predictive power of our factors.

#### 4.10.6 Permission to Reuse and Copyright

Figures, tables, and images will be published under a Creative Commons CC-BY licence and permission must be obtained for use of copyrighted material from other sources (including re-published/adapted/modified/partial figures and images from the internet). It is the responsibility of the authors to acquire the licenses, to follow any citation instructions requested by third-party rights holders, and cover any supplementary charges.

## AUTHOR CONTRIBUTIONS

Conceptualization: TSP, VS, SUK, PK, GP. Formal analysis: TSP, KG, MSM, DN. Investigation: TSP, KG, MSM, DN. Funding acquisition: PK, GP. Resources: KG, MSM, DN, VS. Supervision: SUK, PK, GP. Writing – original draft: TSP, SUK, PK. Writing – review and editing: All authors.

## ACKNOWLEDGMENTS

The authors express their gratitude to everyone who contributed to this project. They would like to thank Nora Maleki and Linus Tiemann for their help in developing the VR city and the Pointing Tasks required for these experiments, and Philipp Spaniol for his 3-dimensional art and implementation of differential loading of levels of details with the agents on this scene.

## DATA AVAILABILITY STATEMENT

The preprocessing and analysis scripts for this project are available in the following GitHub repository. The raw data used in this study can be made available upon request by contacting the corresponding author.

